# Biophysical characterization of the SARS-CoV-2 spike protein binding with the ACE2 receptor and implications for infectivity

**DOI:** 10.1101/2020.03.30.015891

**Authors:** Ratul Chowdhury, Costas D. Maranas

## Abstract

SARS-CoV-2 is a novel highly virulent pathogen which gains entry to human cells by binding with the cell surface receptor – angiotensin converting enzyme (ACE2). We computationally contrasted the binding interactions between human ACE2 and coronavirus spike protein receptor binding domain (RBD) of the 2002 epidemic-causing SARS-CoV-1, SARS-CoV-2, and bat coronavirus RaTG13 using the Rosetta energy function. We find that the RBD of the spike protein of SARS-CoV-2 is highly optimized to achieve very strong binding with human ACE2 (hACE2) which is consistent with its enhanced infectivity. SARS-CoV-2 forms the most stable complex with hACE2 compared to SARS-CoV-1 (23% less stable) or RaTG13 (11% less stable) while occupying the greatest number of residues in the ATR1 binding site. Notably, the SARS-CoV-2 RBD out-competes the angiotensin 2 receptor type I (ATR1) which is the native binding partner of ACE2 by 35% in terms of the calculated binding affinity. Strong binding is mediated through strong electrostatic attachments with every fourth residue on the N-terminus alpha-helix (starting from Ser19 to Asn53) as the turn of the helix makes these residues solvent accessible. By contrasting the spike protein SARS-CoV-2 Rosetta binding energy with ACE2 of different livestock and pet species we find strongest binding with bat ACE2 followed by human, feline, equine, canine and finally chicken. This is consistent with the hypothesis that bats are the viral origin and reservoir species. These results offer a computational explanation for the increased infectivity of SARS-CoV-2 and allude to therapeutic modalities by identifying and rank-ordering the ACE2 residues involved in binding with the virus.

## Introduction

The causative agent of coronavirus disease 2019 (COVID-19) was identified in January 2020 to be a novel *β*-coronavirus of the same subgenus as SARS-CoV-1. SARS-CoV-2 strain has caused a dramatically greater number of infections and fatalities and an effective antiviral treatment and vaccine remains elusive to this day. It has been reported that the first step to viral entry is association between the viral spike RBD and human ACE2 protein^1^. There have been several structural analyses^2,3^ of both SARS-CoV-1 and SARS-CoV-2 binding interactions with human ACE2 (hACE2) but no quantitative assessment of the contribution of different residues in the spike RBD towards tight binding or comparisons with its native receptor ATR1. It has been suggested^2,4^ that viral spike binding to hACE2 prevents ATR1 binding with hACE2 but no quantitative comparisons have been drawn. Experimental and computational investigations have focused on the RBD-hACE2 interaction for SARS-CoV-1^5^ and CoV-2^7^, the role of glycosylated spike residues^8^, and the potential impact of the CoV-2’s furin cleavage site^6^.

In this study, we first assess the molecular interactions between the three spike RBDs with the hACE2 complex. We also provide a comparative analysis of the most important RBD residues from all three viral spike proteins that drive binding with hACE2. Using the Rosetta binding energy function to score interactions, we find that SARS-CoV-2 outcompetes the human ATR1 surface receptor protein to preferentially bind hACE2 by 35% quantified using the Rosetta binding energy function. A recent study^9^ explained interactions between hACE2 and SARS-CoV-1 vs. SARS-CoV-2 RBDs using a homology modeled structure of SARS-CoV-2 RBD and only considering five residues from the spike RBDs. Building on these results, we used an experimentally confirmed atomic scale maps (cryo-EM structures) for the SARS-CoV-1 and CoV-2 RBD in complex with hACE2. Because no experimentally resolved RaTG13-hACE2 complex structure is available, we computationally reconstructed a putative one using flexible protein-protein docking (see Methods). We find that the RBD of SARS-CoV-2 binds hACE2 23% stronger than SARS-CoV-1 and 11% compared to RaTG13 quantified using the Rosetta energy function. Extending this analysis to include non-human ACE2 orthologues, we calculated a descending order of binding strength starting with bats and followed by humans, felines, canines, equines, bovines, and finally poultry. This rank order is consistent with a recent experimental report that finds that mammals especially felines are susceptible to SARS-CoV-2, whereas birds, fish, and reptiles are not^10^.

## Results

### Analysis of human ACE2 in complex with spike RBDs from the three different coronavirus strains

Rosetta-based energy minimization of the hACE2-RBD complexes with RBDs from SARS-CoV-1, SARS-CoV-2, and RaTG13 reveals that SARS-CoV-2 exhibits the strongest Rosetta binding score (−48.312 ± 3.4 kcal/mol). SARS-CoV-1 and RaTG13 Rosetta binding energy scores with hACE2 are – 37.308 ± 2.3 and −43.168 ± 2.1 kcal/mol, respectively. In an uninfected human cell, the ATR1 receptor binds to ACE2 to form a receptor complex. Upon infection, the coronavirus presents the RBD of its spike protein to the human ACE2 forming an electrostatically-driven association between the two. Our results indicate that hACE2 can bind with either human ATR1 or the viral spike (but not both simultaneously) as the binding domains overlap. hACE2 forms hydrophobic and strong electrostatic (including pi-pi, and cation-pi) interactions with the binding domain of ATR1 with a Rosetta binding energy of 31.4 kcal/mol which is 35% less strong than the one with the SARS-CoV-2 RBD. The CoV-2 RBD maximally co-opts these interactions to gain entry via strong non-covalent attachment (see **Figure 1**).

**Figure 1.**
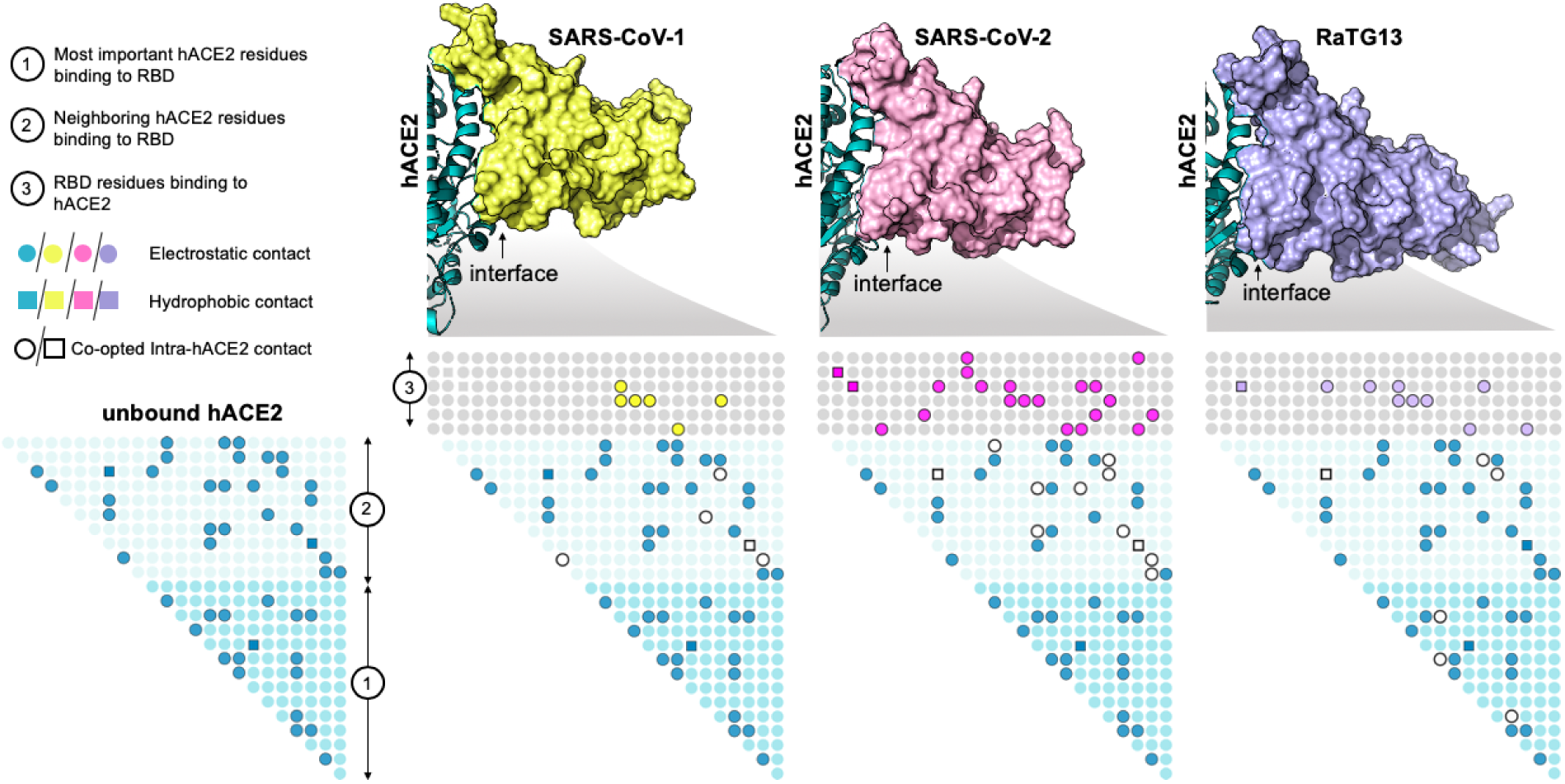
SARS-CoV-2 RBD causes the greatest disruption to the original intra-residue contacts of hACE2 achieving the strongest-binding complex. Shown in the figure are the residue contact maps of the hACE2 receptor in the unbound state and when bound with the viral spike protein RBDs from SARS-CoV-1, SARS-CoV-2, and RaTG13, respectively. Filled dots (in green) represent electrostatic (i.e., circles) or hydrophobic (i.e., squares) intra-residue contacts within hACE2. Open circles and squares in the bound state of hACE2 with RBD signify the lost intra-residue contacts within hACE2 upon binding with the three spikes. Shown in yellow, pink and cyan filled circles and squares are the inter-residues contacts formed upon binding with the three spike RBDs. Filled circles or squares in the light blue region show contacts between hACE2 residues (region 1) that are adjacent to the ones (region 2) contacting the spike RBD (region 3). SARS-CoV-2 disrupts and co-opts the most intra-hACE2 residue contacts forming the most residue contacts between hACE2 and RBD. RBD self-stabilizing contact information and weak (long-range) electrostatic interactions (between 4.5Å and 6.0Å) between the spike and hACE2 are not show in the figure.

To understand the role of the inter-residue interaction network formed during viral entry, we first constructed a contact map depicting all such interactions for the spike-binding interface of unbound hACE2 (see **Figure 1**). We then computed the changes in this contact map upon binding with the RBD of SARS-CoV-1, SARS-CoV-2, and RaTG13. We observe that SARS-CoV-2 more radically co-opts the original contact map of unbound hACE2 to form a highly stabilized hACE2-RBD interface (see **Figure 1**).

We observe that SARS-CoV-2 forms the greatest number of effective hACE2 contacts (11 hydrogen-bonded, eight electrostatic and two hydrophobic) with sixteen RBD residues at the hACE2 binding interface (see **Figure 1**). For example, SARS-CoV-2 RBD residue Phe456 simultaneously forms a hydrophobic contact with hACE2 residue Thr27 (using the side-chain) and an electrostatic stabilization with hACE2 residue Asp30 (using the backbone) (see **Figure 2**). The RaTG13 RBD only forms the hydrophobic interaction whereas the SARS-CoV-1 RBD forms neither (see **Figure 2**). Consequently, a computational alanine scan (see **Figure 3**) reveals that alanine mutation of this position leads to significant loss of hACE2 binding in both SARS-CoV-2 (∼61% reduction) and RaTG13 (∼59% reduction) but not in SARS-CoV-1 (only ∼12% reduction). The spike protein RBD for SARS-CoV-1 (and RaTG13) are only able to form eight (and eleven) strong electrostatic contacts using seven (and ten) RBD residues, respectively. This does not imply that SARS-CoV-1 and RaTG13 only use these residues to bind to hACE2. More than fifteen additional interface residues either form weak electrostatic contacts or are simply non-interacting. **Table 1** lists the hydrogen-bonded interactions between the RBDs and hACE2 along with the corresponding distances. SARS-CoV-2 reforms the original contact map with hACE2 by leveraging 34.1% (15 out of 44) of self-stabilizing contacts around the spike-binding domain to form 21 new complex-stabilizing contacts. SARS-CoV-1 and RaTG13 show weaker attachments as they are able to co-opt only 13.6% and 20.4% contacts, respectively.

**Table 1.**
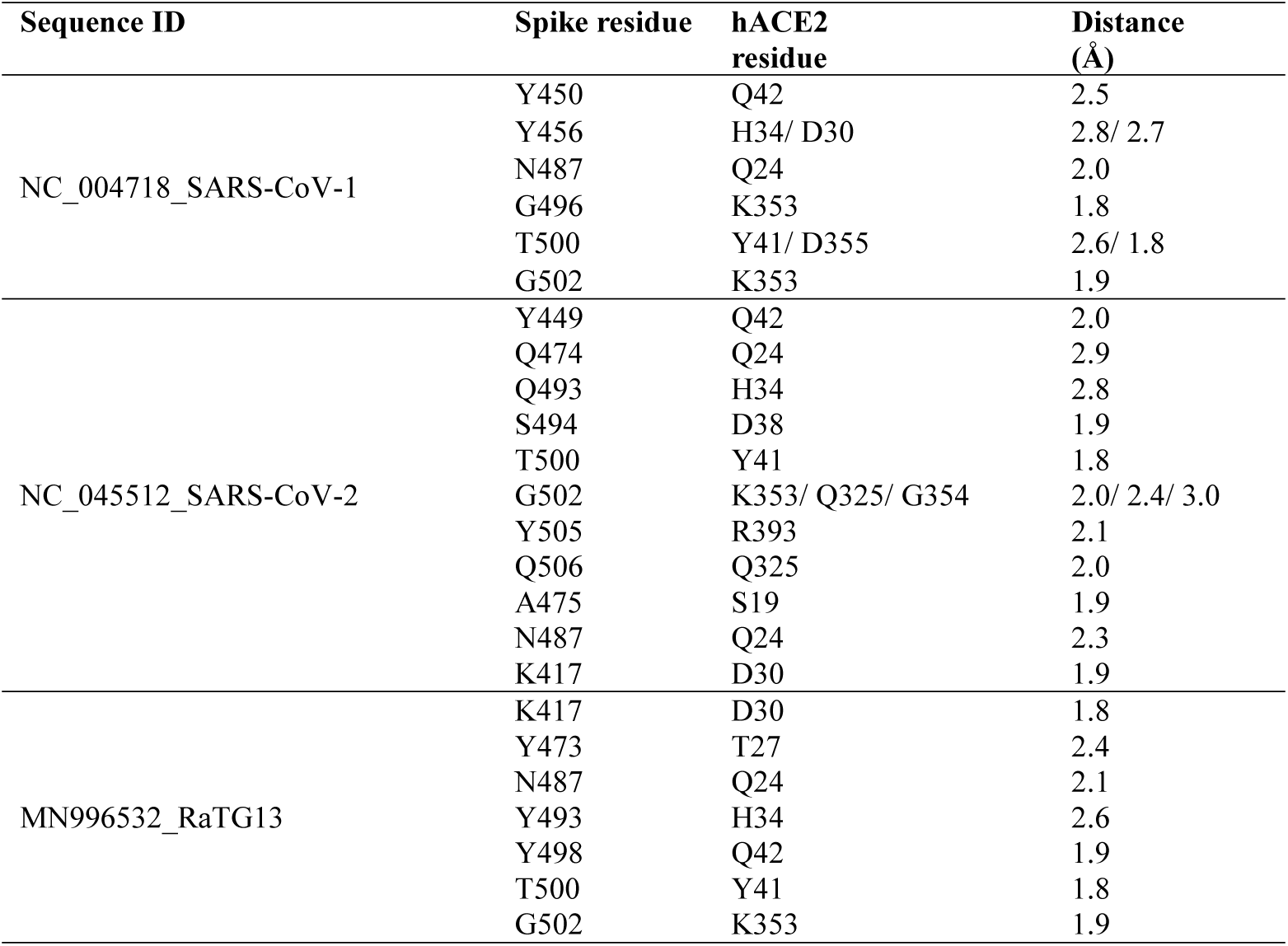
List of hydrogen-bonded contacts between the spike RBDs from (SARS-CoV-1, SARS-CoV-2, and RaTG13) and hACE2.

**Figure 2.**
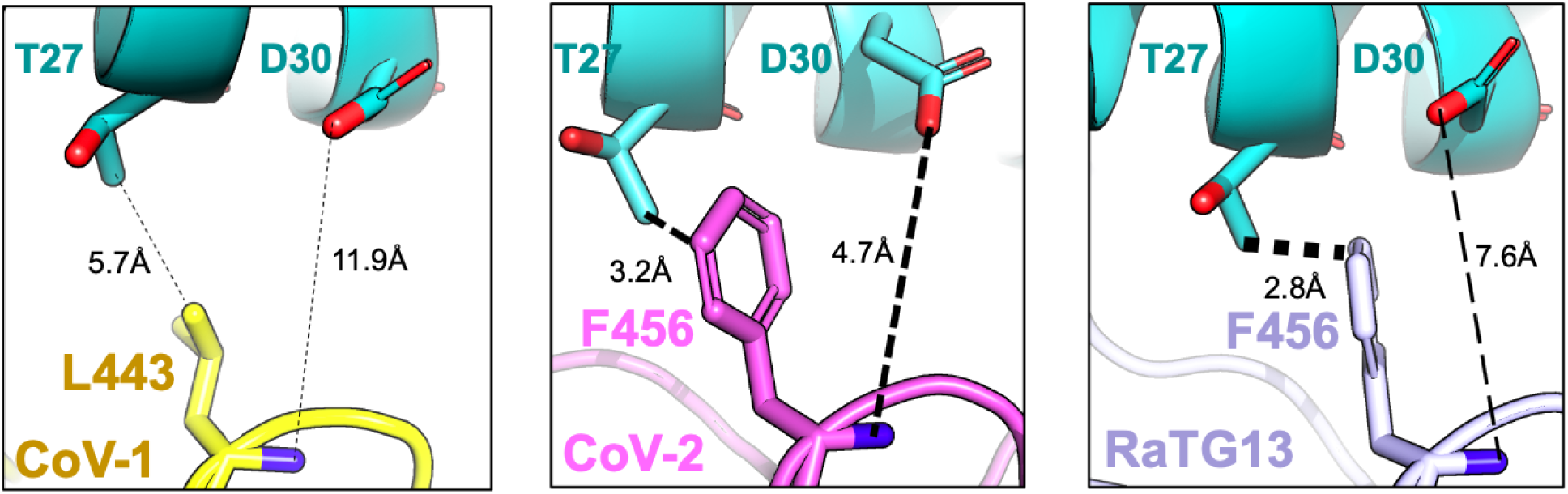
Leu443 present in the SARS-CoV-1 spike RBD is aligned with Phe456 present in SARS-CoV-2 an RaTG13. In SARS-CoV-2, Phe456 simultaneously interacts with hACE2 residues Thr27 and Asp30 whereas only the hydrophobic contact is observed in RatG13. In SARS-CoV-1, Leu443 is unable to establish neither the backbone electrostatic contact nor the hydrophobic stabilization of the methyl group of Thr27 present in hACE2. The thickness of the dashed lines denotes the strength of interaction.

**Figure 3.**
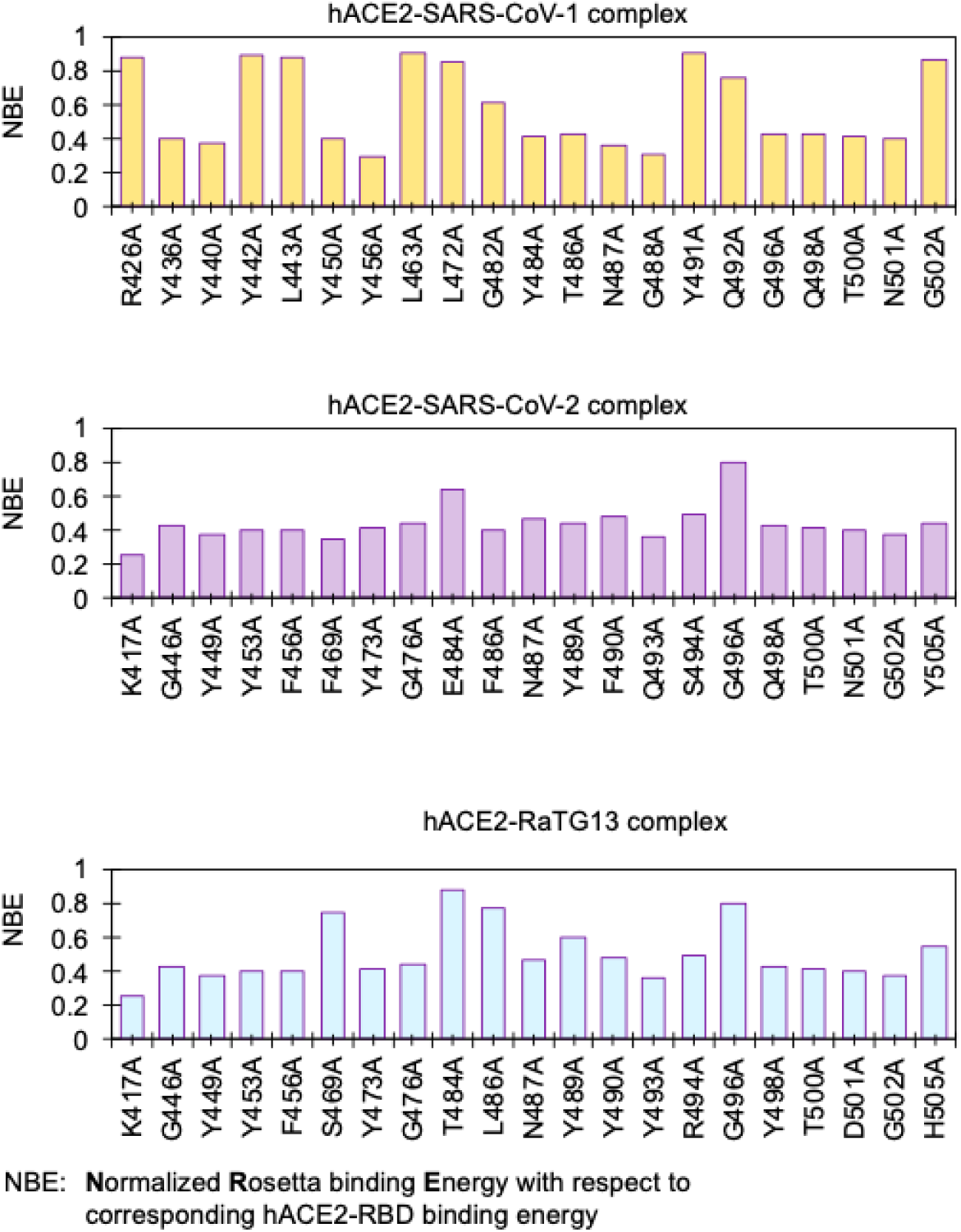
Alanine scan on hACE2 binding residues of spike RBDs of SARS-CoV-2, SARS-CoV-1, and RaTG1 coronavirus. Bars represent the hACE2 Rosetta binding energies upon alanine mutation at the indicated site normalized with respect to binding score prior to mutation. SARS-CoV-2 spike RBD appears to be highly optimized for binding hACE2 as the single mutation to more than 90% of the residues forming the RBD to alanine causes significant reduction in binding energy.

### *In silico* alanine scanning to identify spike residues most important for hACE2 binding

Each one of the hACE2 binding residues from the three viral spike RBDs was computationally mutated to alanine (one at a time) and the resultant hACE2-RBD complexes were energy minimized and scored using the Rosetta energy function. This procedure assesses how important is the identity of the native residues by defaulting them to alanine and observing whether this significantly affects binding. The percent loss of hACE2 binding upon an alanine mutation was used as a proxy score for assessing the importance of each RBD residue in binding and subsequent pathogenesis. The results from the alanine scan study (see **Figure 3**) reveal that ∼90% (19 out of 21) of the hACE2-binding residues of SARS-CoV-2 are important for complex formation. Even a single mutation to alanine of any of these residues lowers the binding score by more than 60%. These results imply that the SARS-CoV-2 RBDs of the spike protein are highly optimized for binding with hACE2. We note that positions Lys417 and Gly502 have one of the strongest impacts on binding (78% and 79% reduction upon mutation to Ala, respectively). This is because they help establish one strong electrostatic contact with Asp30, and three with Gln325, Lys353, and Gly354 (as listed in **Table 1**). The computational alanine scanning results identify the same three residues Phe486, Gln493, and Asn501 to be important for hACE2 binding as proposed by Wan *et al.*^9^. We find that Phe486, Gln493, and Asn501 each establish three new contacts, consequently their mutation to Ala (even for only one of them) leads to loss of ACE2 binding by more than ∼62.5%.

Alanine scanning results of the spike protein RBD of SARS-CoV-1 show less significant penalty to the binding score upon mutation to alanine. Only twelve residues are involved in strong electrostatic coupling with hACE2 residues, out of which six are hydrogen bonded (indicated in **Table 1**). In summary, alanine scans indicate that SARS-CoV-2 has the highest number of “effectively” interacting residues at the ACE2 binding interface whereas the SARS-CoV-1 spike forms only a few strong hACE2 connectors with a large number of “idle” interface residues (43% – 9 out of 21) which do not affect hACE2 binding upon mutation to alanine. RatG13 appears to be between the two with 13 strong electrostatic interactors (61% – 13 out of 21), out of which seven are hydrogen bonded, and only four idle residues at the interface (i.e., residues Thr484, Leu486, Gly496, and Tyr505).

### Presence of tyrosine and glycine residues in the hACE2 binding domains of these spike proteins

All three viral RBDs are enriched in tyrosine residues. As many as 26.3% (5 out of 19 residues) of the SARS-CoV-1 RBD residues, 25% (4 out of 16 residues) for SARS-CoV-2, and 29% (5 out of 17 residues) for RaTG13 are tyrosine residues. We have not explored the phylogenetic basis for the presence of tyrosine residues but they do seem to be important for conferring high binding affinity spike and hACE2 for both SARS-CoV-2 and RaTG13, as alluded to by the alanine scan results (see **Figure 3**). In contrast, the tyrosine residues in SARS-CoV-1 (Tyr442, Tyr475, and Tyr491) only constitute self-stabilizing electrostatic contacts. We use **Figure 4a** to explain one representative case of interface tyrosine residues from all three RBDs: SARS-CoV-1 (Tyr442 and Asn473), SARS-CoV-2 (Tyr473 and Tyr489), and RaTG13 (Tyr473 and Tyr489).

**Figure 4.**
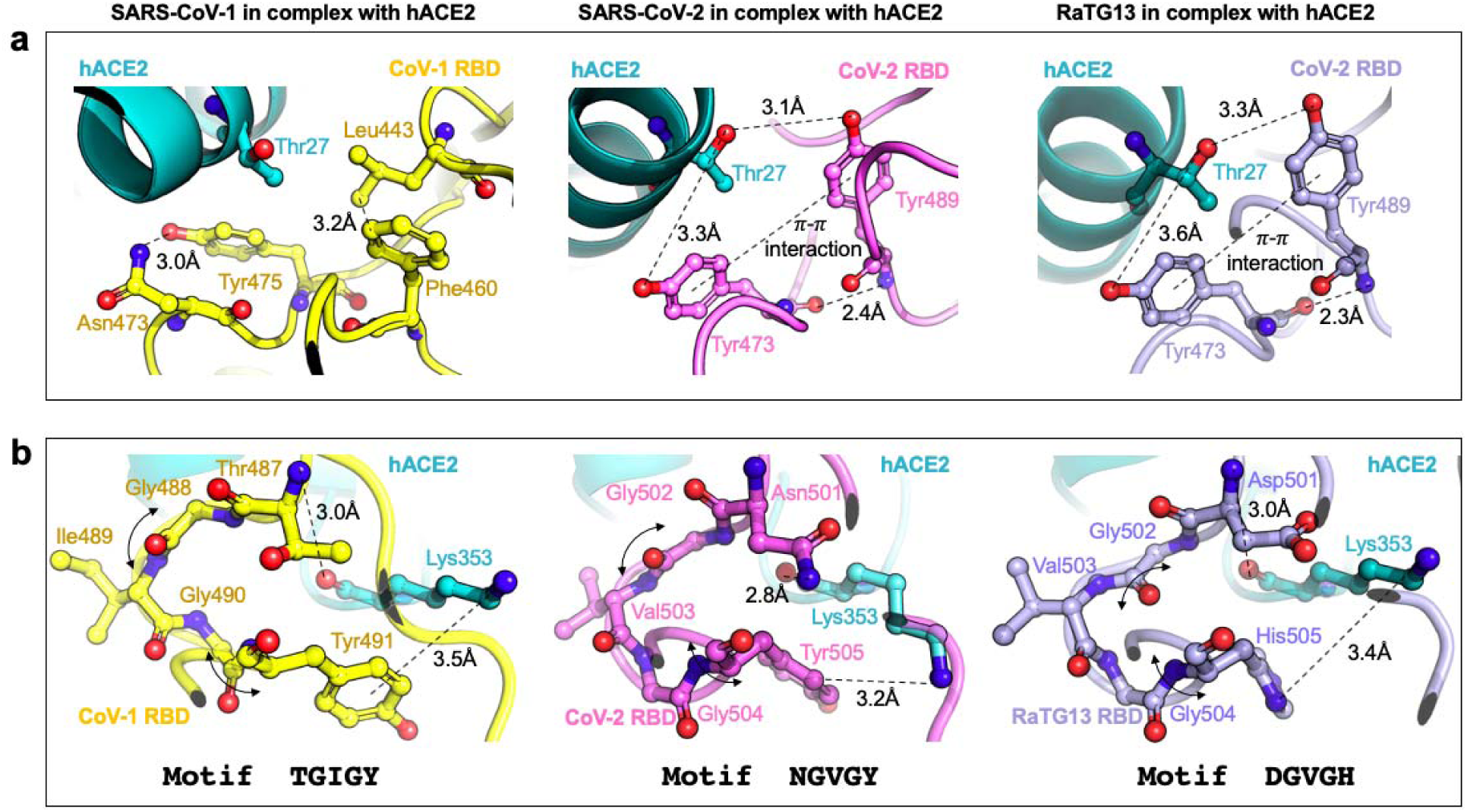
**(a).** The role of tyrosine residues in SARS-CoV-2 and RaTG13 RBD is to form strong contacts with hACE2 residues while in SARS-CoV-1 they are primarily responsible for forming stabilizing contacts within the spike and are hence unavailable for hACE2 binding. **(b)** The role of glycine residues in both all three RBDs is t provide a xGzGx motif for binding hACE2 Lys353 using a strong electrostatic (or cation-interaction). Here, ‘x’ is a polar residue, and ‘z’ a short chain hydrophobic residue (Ile or Val). The glycine residues along with residue ‘z’ offer a hinge to present polar residue ‘x’ for strong electrostatic interactions with hACE2 residue Lys353.

The SARS-CoV-2 and RaTG13 Tyr473 and Tyr489 backbones, even though present in a loop, are mutually stabilized by hydrogen bonding and the side chains are locked in place by a pi-pi aromatic interaction between the phenyl rings. This enables both of these tyrosine side-chains to form a strong electrostatic contact with the Thr27 side-chain of hACE2. It is thus unsurprising that mutation of either Tyr473 or Tyr489 (in both SARS-CoV-2 and RaTG13) to alanine results in a similar (>58%, respectively as shown in **Figure 3**) reduction in binding with hACE2. In contrast, in the energy minimized complex of SARS-CoV-1 RBD with hACE2 both Tyr442 and Tyr475 (see **Figure 4a**) only contribute to internal stability of the spike by forming strong electrostatic contacts with RBD residues Trp476 and Asn473. They are therefore unavailable (or too far > 6.0Å) for binding with the neighboring hACE2 residues.

Next, we focus on the role of glycine residues (see **Figure 4b**) in all three spike RBDs which form important electrostatic contacts with hACE2 as they lead to more than 55% loss of binding (on average) upon mutation to alanine. We chose to study in detail one such representative glycine from all three spike protein RBDs –Gly488 and Gly490 from SARS-CoV-1 and Gly502 and Gly504 from SARS-CoV-2 and RaTG13.

Interestingly, for all three variants the interaction with the hACE2 residue Lys353 with glycine residues in the spike protein is the same. Atomic coordinates of both these complexes were independently, and experimentally confirmed by Song *et al.*^11^ in 2018 and Wang *et al.* in 2020 (manuscript unpublished but structure deposited at – www.rcsb.org/structure/6lzg). Both SARS spike RBDs use a combination of a cation-π and a strong electrostatic interaction to bind with Lys353 whereas RaTG13 uses two electrostatic contacts. One electrostatic interaction is mediated by Thr487 in SARS-CoV-1 and Asn501 (and Asp501) in SARS-CoV-2 (and RaTG13). Two glycine residues and a short hydrophobic residue (‘z’ – Val or Ile) brings Thr487, Asn501, and Asp501 for SARS-CoV-1, SARS-CoV-2, and RaTG13, respectively, within strong electrostatic reach of Lys353 while ensuring another cation-π or an electrostatic interaction between Tyr491, Tyr505, and His505 residues, respectively (see **Figure 4b**). Mutation Y491A for SARS-CoV-1 has no effect on hACE2 binding but Y505A (and H505A) in SARS-CoV-2 (and RaTG13) reduces binding by more than 40%. However, alanine mutation to any of the hinge glycine residues leads to >70% loss of hACE2 binding in all three RBD-hACE2 complexes. Thus, we recover the strong functional motif xGzGx in the spike RBD which is conserved between all three SARS-CoV strains.

Analysis of the three hACE2 binding interfaces (see **Figure 5a-c**) demonstrate that even though all three spike proteins have a similar number of total interface residues (see **Figure 5f**), SARS-CoV-2 establishes more hydrogen bonded contacts (see **Figure 5g)** followed by RaTG13 and SARS-CoV-1. Consequently, SARS-CoV-2 exhibits the strongest Rosetta binding energy with hACE2 (see **Figure 5d**) calculated using ten unique Rosetta energy minimization trajectories. Interestingly, RaTG13 spike residues occupy the largest number of hACE2 residues resulting in the highest reduction (∼14% more than SARS-CoV-2) of solvent accessible surface area (SASA) (see **Figure 5e**). Nevertheless, the associated Rosetta binding energy is 11.2% less than the one for SARS-CoV-2 which forms overall stronger hydrogen-bonded contacts.

**Figure 5.**
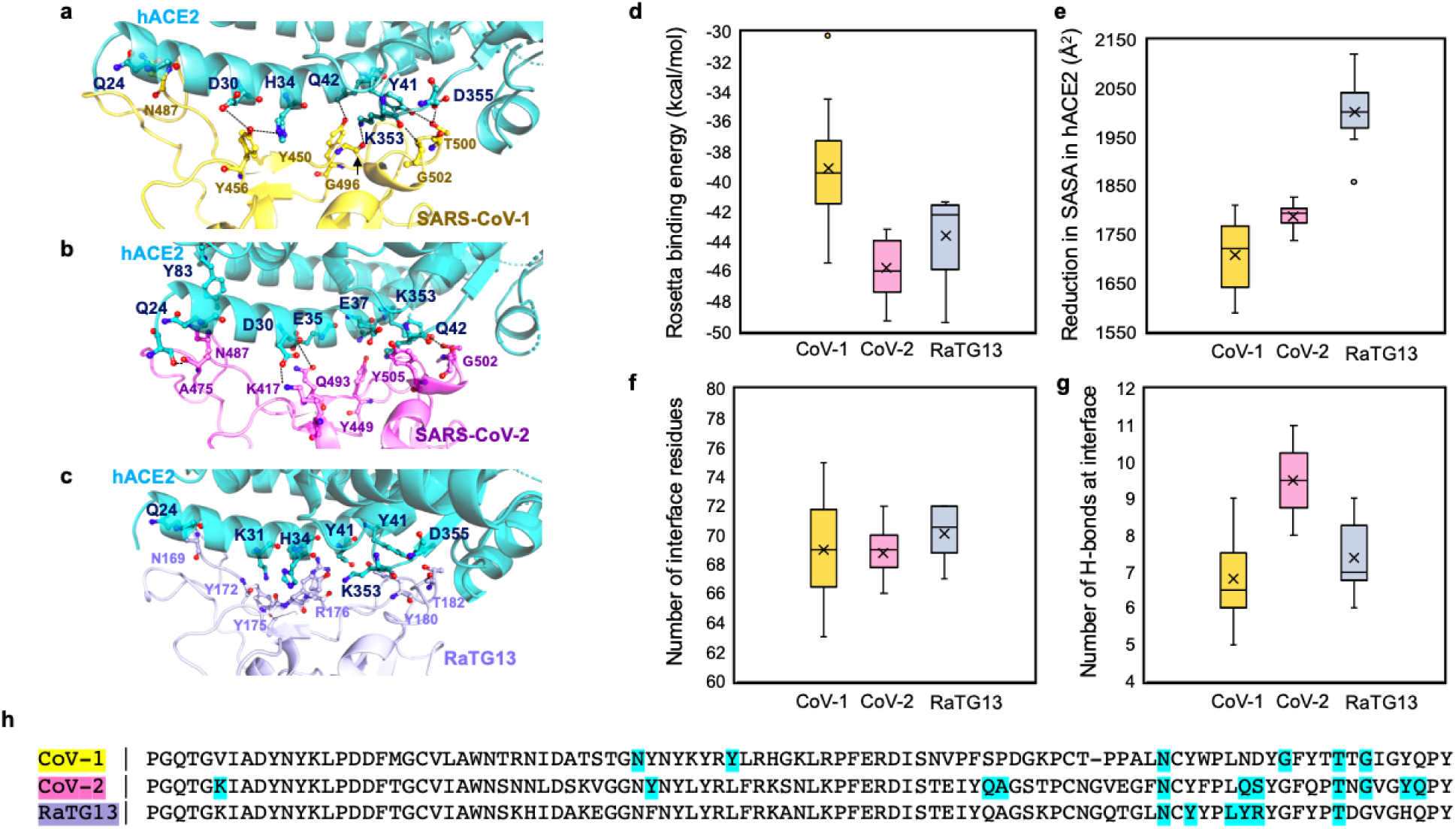
**(a-c)**. hACE2 binding interfaces of the three spike proteins with six hydrogen-bonded contacts from each of them indicated. **(d)** Rosetta binding energies between spike RBD and hACE2 averaged from ten independent binding energy minimization trajectories. **(e)** RaTG13 shows the highest reduction of hACE2 solvent accessible surface area (SASA). **(f-g)** Even though RaTG13 recruits the highest number of interface residues, SARS-CoV-2 forms the most hydrogen-bonded contacts with hACE2. **(h)** The sequence alignment of the three RBDs is shown and the residues establishing hydrogen bonds with hACE2 are highlighted in cyan.

### Competitive hACE2 binding of the spike RBDs and angiotensin receptor (ATR1)

So far, we examined the biophysical characterization of hACE2 binding with the spike protein. However, in an uninfected cell, through the action of the renin angiotensin system (RAS), hACE2 forms a complex with the angiotensin 2 receptor type I (ATR1)^12^. Due to the lack of an experimentally resolved structure for the hACE2-ATR1 complex, we used protein-protein docking and Rosetta binding energy screening to identify the most stable configuration of the complex. Analysis of the hACE2-ATR1 binding interface reveals 41 hACE2 residues and 26 ATR1 residues at the interface connected by five strong electrostatic contacts and several long range weak electrostatic contacts. We find that eleven SARS-CoV-2 RBD binding residues of hACE2 are shared by the ATR1 binding region. Moreover, the SARS-CoV-2 spike protein binds hACE2 with ∼35% better binding score than ATR1 binds hACE2. RaTG13 and SARS-CoV-1 exhibit ∼21% and ∼5% better Rosetta binding energies, respectively with hACE2 compared to the hACE2-ATR1 complex. They also share only nine and eight residues, respectively with the ATR1 binding interface of hACE2 as opposed to eleven for SARS-Cov-2 (see **Figure 6**). Rosetta binding calculations therefore suggest that SARS-CoV-2 can more effectively than CoV-1 outcompete the hACE2-ATR1 complex thus possibly facilitating the formation of the hACE2-spike complex. This is in line with the respective Cov-1 vs. Cov-2 infectivities.

**Figure 6.**
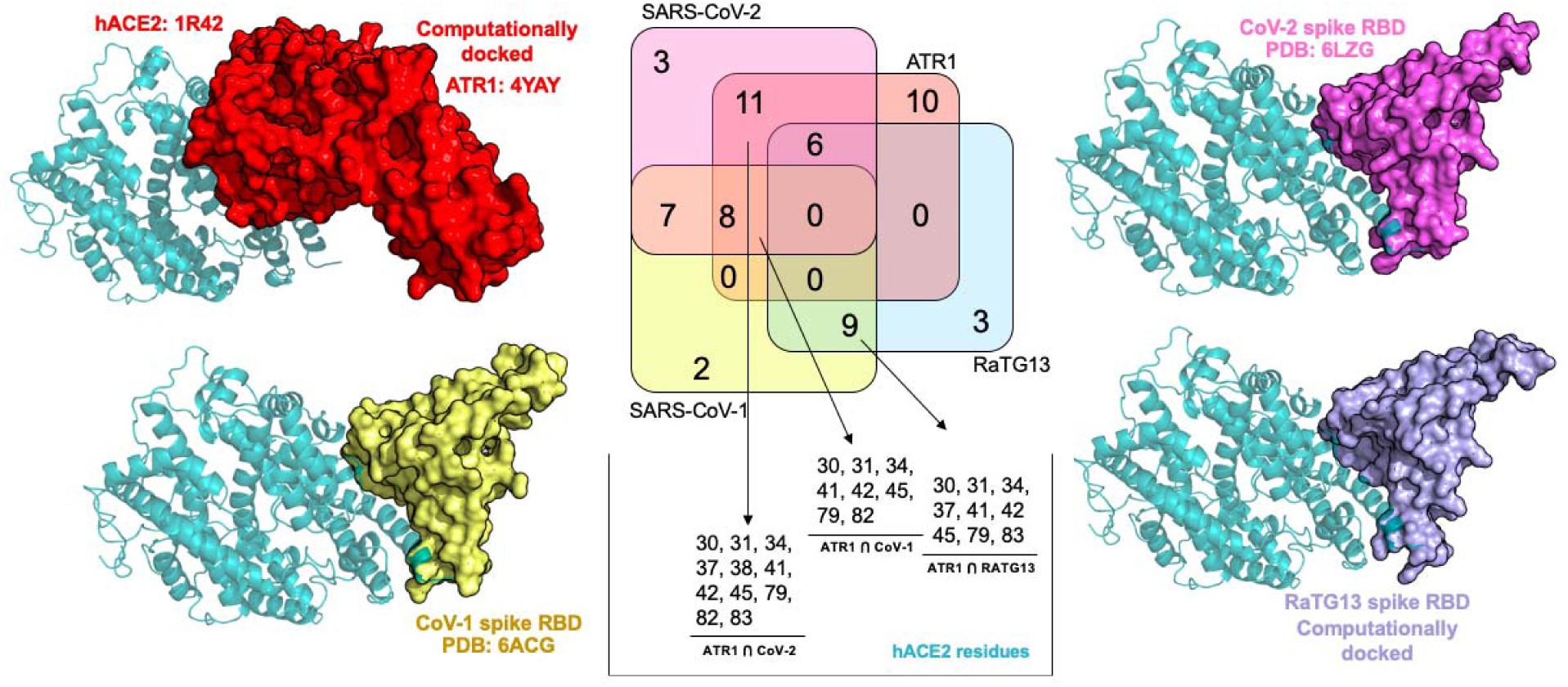
hACE2 complexes with ATR1, SARS-CoV-1, SARS-CoV-2, and RaTG13 spike RBDs along with the number of shared hACE2 residues (Venn diagram) at their respective binding regions is shown. Residue positions that are shared between ATR1 and the three spike RBDs (SARS-CoV-1, SARS-CoV-2, and RaTG13) have been listed.

We computationally explored the potentially available margin of improvement for the binding affinity of SARS-CoV-2 with hACE2 using the IPRO^13^ protein design software. We allowed all 21 contacting residues of the RBD of the spike protein to simultaneously mutate. We run two separate design trajectories and, in both cases the best design achieved an approximately 23% improvement in binding affinity using the Rosetta scoring function. This improvement is less than the difference between the calculated binding scores of SARS-CoV-1 and SARS-CoV-2 implying that SARS-CoV-2 has already achieved most of the theoretically possible binding affinity gain with hACE2 compared to SARS-CoV-1. Interestingly, the network of glycine residues in SARS-CoV-2 is conserved in all redesigned RBDs.

A recent report^14^ analyzes that humans can transfer SARS-CoV-2 to domesticated animals such as dogs, cats, ducks, and chickens in varying degrees. However, animal-to-human transmission has not been observed^15^. Similar to SARS-CoV-1^16^, felines are more susceptible to SARS-CoV-2 followed by canines^17^ whereas chickens and ferrets are less susceptible^17^. The calculated Rosetta binding energies do not follow the trends (R^2^=0.383) expected from simply their respective sequence identities with the human ACE2. Interestingly, even though the ACE2 (Uniprot Entry: G1PKW9_MYOLU) of the little brown bat (*Myotis lucifugus*) is quite different from human (similarity 84.5%, identity 66.7%), we predict a stronger Rosetta binding energy (by about ∼5.6%). This is due to the formation of nine electrostatic contacts and one pi-pi stacking. Strong binding with bat ACE2 may be a consequence of the SARS-CoV-2 origins. In all other cases, the Rosetta binding energies of ACE2 with the spike protein were at most 78.3% of the one calculated with hACE2. We found that feline ACE2 had the closest (78.3% of hACE2-CoV-2) Rosetta binding energy with the spike compared to other pet or livestock animals.

## Discussion

In this effort we apply Rosetta binding analysis to gain insight onto possible biophysical factors that may explain the difference in pathogenicity of SARS-CoV-2 in comparison to SARS-CoV-1 and RaTG13. Multiple lines of computational evidence indicate that the spike RBD binds hACE2 through electrostatic attachment with every fourth residue on the N-terminal alpha-helix (starting from Ser19 to Asn53) as the turn of the helix makes these residues solvent accessible. Results from computational models of canine, feline, bovine, equine, and chicken ACE2 in complex with SARS-CoV-2 spike RBD recapitulates infectivity potential observed so far and pinpoint bat ACE2 as the most highly optimized for binding the SARS-CoV-2 spike protein.

## Methods

We have used experimentally determined coordinates of SARS-CoV-1 and SARS-CoV-2 in complex with ACE2 (PDB accessions: 6ACG^11^ and 6LZG – www.rcsb.org/structure/6lzg, respectively). RaTG13 RBD model was built using the iTasser program^18^. Similarly, unbound ATR1 structure (PDB: 4YAY^19^) was also separately downloaded and docked against hACE2 using protein-protein docking scripts from Z-DOCK 3.0^20^. ZDOCK uses pairwise shape-complementarity, electrostatics, and implicit solvation terms in scoring the docked poses. Implicit solvation treats the water as a dielectric continuum. The rotational sampling interval was set to 10°. Clustering of the docked poses were done at an 8 Å cutoff. Subsequently, PyRosetta^21^ scripts were written to rank and identify the most stable complexes from each cluster which were then energy-minimized and re-ranked. Finally, the complex which ranked high in stability and binding scores was chosen as the model. An alanine scan was again performed using PyRosetta scripts, where the computational models of the alanine variants were first generated, energy minimized, and hACE2 binding scores computed. The hACE2 interface definitions for each binding partner (RBDs and ATR1) were obtained by feeding the energy minimized protein-protein complexes through the *find_contacts* module of OptMAVEn-2.0^22^.

We used the three-dimensional atomic coordinates of the experimentally determined human ACE2 (hACE2) in complex with SARS-CoV-2 spike RBD (PDB id: 6ZLG https://www.rcsb.org/structure/6lzg) as a backbone template to repackage the updated residue side-chains of bat, feline, canine, bovine, equine, and chicken ACE2. A python script was prepared to execute multiple times the iTasser program^18^. First, a fragment structure assembly was performed using replica-exchange Monte Carlo^23^ followed by clustering of decoy ACE2 structures generated using the SPICKER protocol^24^. Finally atomic-level backbone and side chain refinement was performed using fragment-guided molecular dynamics simulations (FG-MD)^25^ for 50ns for each structure. All five ACE2s were subsequently docked with the SARS-CoV-2 spike RBD protein whose 3D coordinates were downloaded from the hACE2-spike complex (PDB id: 6LZG).

## Author Contributions

RC, and CDM conceived, designed, and wrote the study.

## Acknowledgement

RC thanks Debolina Sarkar for advice on the renin angiotensin system and also editing the paper. This activity was partially enabled by research conducted within the Center for Bioenergy Innovation (DE-SC0018420) and NSF CBET1703274. All simulations were performed on the Institute for Computational and Data Sciences Advanced CyberInfrastructure (ICDS-ACI) high-performance computing (HPC) facility at the Pennsylvania State University.

